# Let it bud: an ultrastructural study of *Cryptococcus neoformans* surface during budding events

**DOI:** 10.1101/2020.05.15.098913

**Authors:** Glauber R. de S. Araújo, Carolina de L. Alcantara, Noêmia Rodrigues, Wanderley de Souza, Bruno Pontes, Susana Frases

## Abstract

*Cryptococcus neoformans* is a fungal pathogen that causes life-threatening infections in immunocompromised individuals. It is surrounded by three concentric structures that separate the cell from the extracellular space: the plasma membrane, the cell wall and the polysaccharide capsule. Although several studies have revealed the chemical composition of these structures, little is known about their ultrastructural organization and remodeling during *C. neoformans* budding event. Here, by combining the state-of-the-art in light and electron microscopy techniques we describe the morphological remodeling that occurs synergistically among the capsule, cell wall and plasma membrane during budding in *C. neoformans*. Our results show that the cell wall deforms to generate a specialized budding region at one of the cell’s poles. This region subsequently begins to break into layers that are slightly separated from each other and with thick tips. We also observe a reduction in density of the capsular polysaccharide around these specialized regions. Daughter cells present a distinct spatial organization, with polysaccharide fibers aligned in the direction of budding. In addition, to control the continuous openings between mother and daughter cells, the latter developed a strategy to shield themselves by forming multilamellar membrane structures in conjunction with their capsules. Together, our findings provide compelling ultrastructural evidence for a dynamic *C. neoformans* surface remodeling during budding and may have important implications for future studies exploring these remodeled specialized regions as drug-targets against cryptococcosis.

## 1. Introduction

Fungal infections that cause systemic mycoses have become a major threat, a clinical and a pharmaceutical challenge since the end of the 20^th^ century, especially affecting individuals with an immunological impairment (Perfect, 2013). There are evidences showing that the increase in fungal infections might be correlated with glucocorticoid therapy, immunotherapy, oncological and hematological diseases, increased number of transplants, surgical procedures, individuals living with acquired immunodeficiency syndrome (AIDS), among others (Henao-Martínez and Beckham, 2015; Liao et al., 2016; Singh et al., 2008).

*Cryptococcus* spp., of which *Cryptococcus neoformans* is the main representative of the genus, is a basidiomycete that presents itself as a haploid and spherical yeast surrounded by a polysaccharide (PS) capsule, a unique feature among eukaryotes (D. McFadden et al., 2006). These cells commonly have an average diameter ranging from 2 to 8 μm; however, under certain conditions of physical and/or chemical stresses, they can reach up to 100 μm in diameter, the so-called “titan cells” (Faganello et al., 2006; Trevijano-Contador et al., 2018; Zaragoza, 2019; Zaragoza et al., 2003). *Cryptococcus* spp. has a global distribution and causes about 625,000 deaths per year worldwide (Park et al., 2009). The host becomes infected after inhaling spores or desiccated yeasts (Ellis and Pfeiffer, 1990) and the infection can either take its latent form, without causing any clinical symptoms, or manifest itself in the acute form of the disease (Goldman et al., 2010). Given that *Cryptococcus* spp. has a special tropism toward the Central Nervous System (CNS) and can colonize the CNS through many concomitant infection routes (Mitchell et al., 1995), one can consider cryptococcal meningitis as the most severe cryptococcosis scenario (Casadevall and Perfect, 2008).

The success of the infection is based on the ability of the fungus to evade the host’s immune system. During its evolution, *Cryptococcus* spp. developed several adaptation mechanisms, known as virulence factors. Some examples are: **(I)** melanin production and cell wall remodeling (resistance to cell-mediated death and immunomodulation) (Doering et al., 1999; Gómez and Nosanchuk, 2003; Huffnagle et al., 1995; Liu et al., 1999; Wang et al., 1995), **(II)** production of superoxide dismutase (protection against toxic free radicals) (Cox et al., 2003) **(III)** phospholipase and urease secretion (intracellular growth, diffusion and proliferation) (Cox et al., 2001, 2000) **(IV)** phenotypic switching (immune evasion) (Fries et al., 2001; Goldman et al., 1998), **(V)** cellular gigantism (immune evasion) (Okagaki et al., 2010; Trevijano-Contador et al.,2018; Zaragoza, 2019; Zaragoza and Nielsen, 2013) and **(VI)** PS production, which is the main virulence factor used by *C. neoformans* (Araujo et al., 2012; Zaragoza, 2019; Zaragoza et al., 2009). Most of these features are believed to have been acquired through selective pressures and are likely to be the result of interactions with environmental predators, such as amoebae and nematodes (Albuquerque et al., 2019; Casadevall and Pirofski, 2007).

After production, *Cryptococcus* spp. PS can be either secreted to the extracellular milieu through vesicles (Rodrigues et al., 2007) or transported to the cell wall where it forms the physical structure of the capsule *in situ*. Depending on its fate, PS acquires different physicochemical and rheological properties (Araújo et al., 2019; Pontes and Frases, 2015; Zaragoza, 2019). Due to its unique morphology and the fact that it is pivotal for the establishment of pathogenesis, the PS capsule is the most distinctive feature of the *Cryptococcus* genus. It is highly dynamic, extremely hydrophilic and can be modified in response to the environment. This structure appears at the surface of the cell wall and its main roles are to protect the cell against host’s defense factors and to interfere with immune response mechanisms (Perfect and Casadevall, 2011). In order to anchor to the cell wall, the PS molecules from the capsule interact with α-1,3 glucans (Reese and Doering, 2003). However, the full mechanism that dictates the interaction between capsule and cell wall is far from being completely understood but is thought to involve molecular interactions between the components of both structures.

The fungal cell wall is an intricate network of macromolecules such as lipids, proteins and other polymers like glucans, mannans, galactomannans and chitin. This structure is considered to be a primary determinant of the fungi resistance to stress and environmental aggressions. It provides not only strength and rigidity to maintain the cell conformation but also flexibility to support morphological changes, such as cell growth and budding (Adams, 2004; Roncero, 2002; Ruiz-Herrera et al., 2002). The cryptococcal cell wall also serves as the scaffold for the assembly/anchoring of the PS capsule. Genetic interruptions of its synthesis reduce cell viability and decrease capsule assembly, often producing avirulent mutants. These features make the capsule an attractive target for the development of antifungal therapies, especially because mammalian cells do not have equivalent structures (Wang et al., 2018).

The cell wall is comprised by a matrix containing glycoproteins and glucose (Glc), *A*-acetylglycosamine (GlcNAc) and glucosamine (GlcN) polymers, whose main constituents are glucans, chitin and chitosans (Perfect and Casadevall, 2011). The glucans are divided into α- and β-glucans. A large fraction of α-glucans present α-1,3 links (Bose et al., 2003; James et al., 1990; Wang et al., 2018) whereas the majority of β-glucans are comprised by β-1,3 and β-1,6 bonds (James et al., 1990; Manners et al., 1973; Wang et al., 2018). Chitin, another constituent of the cell wall, is a water-insoluble β-1,4-GlcNAc polymer, that associates with one another to form chitooligomers (chitooligosaccharides). These chitooligomers contain between three to twenty residues of β-1,4-GlcNac, which provides the cell wall with rigidity and structural integrity under various environmental conditions. In *C. neoformans*, chitooligomers are also incorporated into the capsular network and interact with glucuronoxylomannan (GXM) to form complex glycans. Chitin-derived oligomers have also been shown to regulate capsular architecture in *C. neoformans* cells, playing an indirect role in cryptococcal pathogenesis. Finally, they were also detected at the capsular surface, suggesting their potential to be recognized by host receptors, possibly affecting cryptococcal pathogenesis (Fonseca et al., 2013). Cell wall chitins can also be deacetylated to generate chitosan, a more soluble and flexible glucosamine polymer. *C. neoformans* have high levels of chitosan that can exceed chitin amounts up to 10 times (Banks et al., 2005). Cells without chitosan grow slower than the wild type and present impaired cell integrity and reduced virulence in animal models (Baker et al., 2011). Overall, glycoproteins are crucial components of the cell wall in fungi, as they act in critical processes, including signal transduction, conjugation, cell wall synthesis and iron acquisition. These proteins are modified by *N*-oligosaccharide and *O*-oligosaccharide bonds, usually mannosylated structures, which syntheses are initiated in the endoplasmic reticulum and in the Golgi complex (Wang et al., 2018). Glycans linked to the cryptococcal proteins contain xylose (Xyl) and Xyl-phosphate moieties (Lee et al., 2015; Park et al., 2012; Reilly et al., 2011). Even though the full spectrum of glycans has not yet been completely elucidated, it is known that it contains sialic acid that plays an anti-phagocytic role and may represent a virulence factor in the initial stages of infection (Rodrigues et al., 2002).

Since the last century there has been a great interest in deciphering the chemical composition of *C. neoformans* cell surface; however, little is known about its ultrastructural organization and remodeling during important events of *C. neoformans* biology. In the present work, we combined the most up-to-date light and electron microscopy techniques to describe the morphological remodeling that synergistically occurs between the capsule, the cell wall and the plasma membrane during the budding phenomenon in *C. neoformans*.

## 2. Materials and Methods

### 2.1 Microorganisms

The strain used in this work was *C. neoformans* var. *grubii* H99 (clinical isolate, kindly provided by Professor Arturo Casadevall - Johns Hopkins Bloomberg School of Public Health, Baltimore, Maryland, USA), a wild type strain available in the American Type Culture Collection (ATCC catalog number 208821).

### 2.2 Capsule induction and culture conditions

Yeasts were grown in Sabouraud Dextrose Broth (Kasvi, PR, Brazil) medium at 37°C with constant agitation at various times, depending on the experimental conditions. For video microscopy observations, the yeasts were taken directly from Sabouraud Dextrose Agar (Kasvi, PR, Brazil) and, subsequently, added in liquid culture medium and processed, as described below. In order to induce capsule formation, the yeasts were kept at 37°C for 7 days in a nutrient-deprived medium called Minimal Medium (MM) containing only 15 mM glucose, 10 mM MgSO_4_7.H_2_O, 29 mM KH_2_PO_4_, 13 mM glycine and 3 μM thiamine (all compounds from Merck Millipore, Darmstadt, Germany).

### 2.3 Video microscopy

An initial inoculum of 10^4^ cells/mL in Sabouraud medium was added to 35 mm glass bottom dishes (Thermo Scientific ™ Nunc Glass Bottom Dish, Waltham, MA, USA) and observed under a Nikon Eclipse TE300 inverted microscope equipped with a CFI Achromatic LWD ADL 40X objective lens. For 5 hours, phase contrast images were captured every minute using a Hamamatsu C2400 CCD camera (Hamamatsu, Japan). Images were then mounted into stacks and analyzed using the ImageJ 1.8.0 software (NIH, Bethesda, MD, USA - https://imagej.nih.gov/ij/) (Abràmoff et al., 2004; Schneider et al., 2012).

### 2.4 Conventional fluorescence microscopy and structured illumination microscopy (SIM)

Yeast cells (10^6^) were centrifuged at 6,708 *g* for 5 minutes, resuspended in 4% (v/v) paraformaldehyde (Electron Microscopy Sciences, Hatfield, PA, USA) in phosphate buffered saline (PBS) (137 mM NaCl, 2.7 mM KCl, 10 mM Na_2_HPO_4_ and 1.8 mM KH_2_PO_4_) pH 7.2 and incubated for 30 minutes at room temperature. Next, fixed cells were washed twice with PBS and incubated with 1% bovine serum albumin (Sigma Aldrich, Darmstadt, Germany) in PBS for 1 hour at room temperature. The cells were then incubated for another hour at room temperature with 18B7 mAb (10 μg/mL). The 18B7 mAb is a mouse IgG1 with high affinity for GXM from different cryptococcal serotypes (Goldman et al., 1998). After three washes in PBS, cells were incubated with 10 μL/mL of the anti-mouse Alexa Fluor^®^ 594 secondary antibody (Thermo Fisher Scientific, Waltham, Massachusetts, USA) for 1 hour at room temperature. Again, cells were washed with PBS buffer and incubated with Uvitex2B (Polyscience Inc, Warrington, PA, USA) for 1 hour at room temperature and subsequently, vigorously washed 4 times with PBS buffer in order to remove all Uvitex2B dye to minimize background.

Cell suspensions were mounted on glass coverslips and observed using an Axio Observe or an Elyra PS.1 microscope (Zeiss, Germany). Images were acquired with their respective software packages and subsequently processed using ImageJ 1.8.0 software (NIH, Bethesda, MD, USA - https://imagej.nih.gov/ij/) (Abràmoff et al., 2004; Schneider et al., 2012).

### 2.5 Conventional scanning electron microscopy (CSEM)

The cells of interest were washed three times in PBS pH 7.2 and fixed in 2.5% glutaraldehyde solution grade I (Electron Microscopy Sciences, Hatfield, PA, USA) in sodium cacodylate buffer 0.1 M pH 7.2 for 1 hour at room temperature. Then, the cells were washed three times in 0.1 M sodium cacodylate buffer pH 7.2 containing 0.2 M sucrose and 2 mM MgCl2 (Merck Millipore Darmstadt, Germany), and adhered to 12 mm diameter round glass coverslips (Paul Marienfeld GmbH & Co. KG, Germany) previously coated with 0.01% poly-L-lysine (Sigma-Aldrich, Darmstadt, Germany) for 20 minutes. Adhered cells were then gradually dehydrated in an ethanol (Merck Millipore, Darmstadt, Germany) series (30, 50 and 70% for 5 minutes and 95% and 100% twice for 10 minutes). The dehydration procedure was meticulously monitored to prevent PS collapse during air drying or PS extraction due to excessive incubation. The coverslips were then critical-point-dried using an EM DPC 300 critical point drier (Leica, Germany) and mounted on specimen stubs using a conductive carbon adhesive (Pelco Tabs™, Stansted, Essex, UK). Next, the samples were coated with a thin layer of gold or gold-palladium (10-15 nm) using the sputter method (Balzers Union FL −9496, Balzers, FL). Finally, samples were visualized in a scanning electron microscope (Zeiss Evo 10 or FEI Quanta 250) operating at 10-20 kV with an average working distance of 10 mm and images were collected with their respective software packages.

### 2.6 High resolution scanning electron microscopy (HRSEM)

The cells were processed following the same methodology described above (please, see CSEM). However, in order to proceed with HRSEM, the samples were sputtered with a 3 nm thick platinum layer on their surfaces and observed in high resolution electron microscopes, either FEI Magellan™ (FEI Company, Oregon, USA) or Zeiss Auriga-40 (Zeiss, Germany) operating at 1 kV with an average working distance of 2 mm and images were collected using their respective software packages.

Quantification of PS fiber anisotropy was performed using FibrilTool (Boudaoud et al., 2014), an ImageJ plug-in that determines the average orientation of a fiber array. The anisotropy value ranges from a maximum of 1, when all fibers point to the same direction, to a minimum of 0, when fibers are randomly oriented.

### 2.7 Transmission electron microscopy (TEM)

The cells were washed three times in PBS pH 7.2 and subsequently fixed in 2.5% (v/v) glutaraldehyde solution grade I in 0.1 M sodium cacodylate buffer pH 7.2 and microwaved (350 W, 3 pulses of 30 seconds each with an interval of 60 seconds between pulses) (Benchimol et al., 1993; Giberson et al., 2003). Subsequently, the cells were washed three times in 0.1 M sodium cacodylate buffer pH 7.2. The cells were then post-fixed using an Osmium-Thiocarbohydrazide-Osmium (OTO) protocol. Briefly, cells were incubated in a post-fixative 1% (v/v) osmium tetroxide (OsO_4_), 0.8% (v/v) potassium ferrocyanide and 5 mM calcium chloride, in 0.1 M cacodylate buffer (pH 7.2) for 10 min, washed twice in water, and then incubated in 1% (w/v) thiocarbohydrazide (TCH, Sigma, Darmstadt, Germany) in water, for 5 min (Murakami et al., 1983; Seligman et al., 1966; Willingham and Rutherford, 1984). After three washes in water, cells were again incubated in the post-fixative osmium solution for 2 min and finally washed three times in water. Next, cells were gradually dehydrated in an acetone (Merck Millipore, Darmstadt, Germany) series: 50%, 70%, 90% and two subsequent 100%. All the dehydration procedures were performed in the microwave (350 W, 10 seconds pulses for each step). The Spurr resin (Electron Microscopy Sciences, Hatfield, PA) was gradually used to substitute acetone in the following proportions acetone: Spurr (v:v): 3:1, 2:1, 1:1, 1:2, 1:3 and finally pure Spurr. Each mixture was also submitted to the same microwave cycles for 2 minutes except the last step (pure Spurr) that was performed without any radiation. The polymerization step was carried out for 48 hours in an oven at 70°C. The samples were sliced in 75 nm sections under a Leica EM UC7 ultramicrotome (Leica, Wetzlar, Germany), collected onto formvar-coated copper slot grids and submitted to an incubation with 5% (w/v) uranyl acetate in water for 20 minutes and lead citrate for 5 minutes for contrasting. Finally, the samples were observed in a Tecnai™ Spirit microscope operated at 120 kV (FEI Company, Oregon, USA) and images were collected using the microscope software.

### 2.8 Electron tomography and three-dimensional reconstruction

The samples were processed following the same procedures described above (see TEM). However, for electron tomography, a few different steps were performed, as follows. Samples were sliced into 200 nm thick serial sections under a Leica EM UC7 ultramicrotome (Leica, Wetzlar, Germany), collected onto formvar-coated copper slot grids and stained with 5% (w/v) uranyl acetate for 3 minutes and Reynolds’ Lead citrate for 5 minutes. The tomographic series were acquired with an inclination of ± 65° and 1° increments under a Tecnai Spirit™ (FEI Company, Oregon, USA) transmission electron microscope operating at 120 kV with a 2,048 × 2,048 pixels’ matrix CCD camera. Serial tilt series were aligned using Etomo, an open-source software from IMOD package, a set of image processing, modeling and display programs used for tomographic reconstruction and for 3D reconstruction of EM serial sections (Kremer et al., 1996; Mastronarde, 1997). Generated tomograms were reconstructed using 3dmod.

### 2.9 Statistical analysis

Statistical analysis were performed using GraphPad Prism 8.4.0 (GraphPad Software, La Jolla, CA). Student’s *t*-test was used for comparisons.

## 3. Results

### 3.1 *Cryptococcus neoformans* differentiates a region of its cell wall to generate daughter cells

*C. neoformans* divides through budding of daughter cells from mother cells (Lin et al., 2014). To revisit and better characterize this phenomenon, yeast cell cultures were grown in Sabouraud Dextrose Broth medium and were allowed to attach onto coverslips. Next, we acquired phase contrast images of the same field of view every minute for 5 hours. Images were mounted in stacks allowing us to follow the proliferative behavior of the cells. Mother cells (G1) generated their daughters (G1.1) at an average rate of one cell every (1.3 ± 0.3) h (Figure 1 and Supplementary Video 1). Moreover, successive daughter cells always bud from the same regions of their respective mothers, not only from cells that were attached since the beginning (Figures 1A, 1B and 1C and Supplementary Video 1), but also from daughter cells that subsequently attached to the coverslips after budding and also started to generate daughter cells of their own (Figures 1C and 1D and Supplementary Video 1).

**Figure 1:**
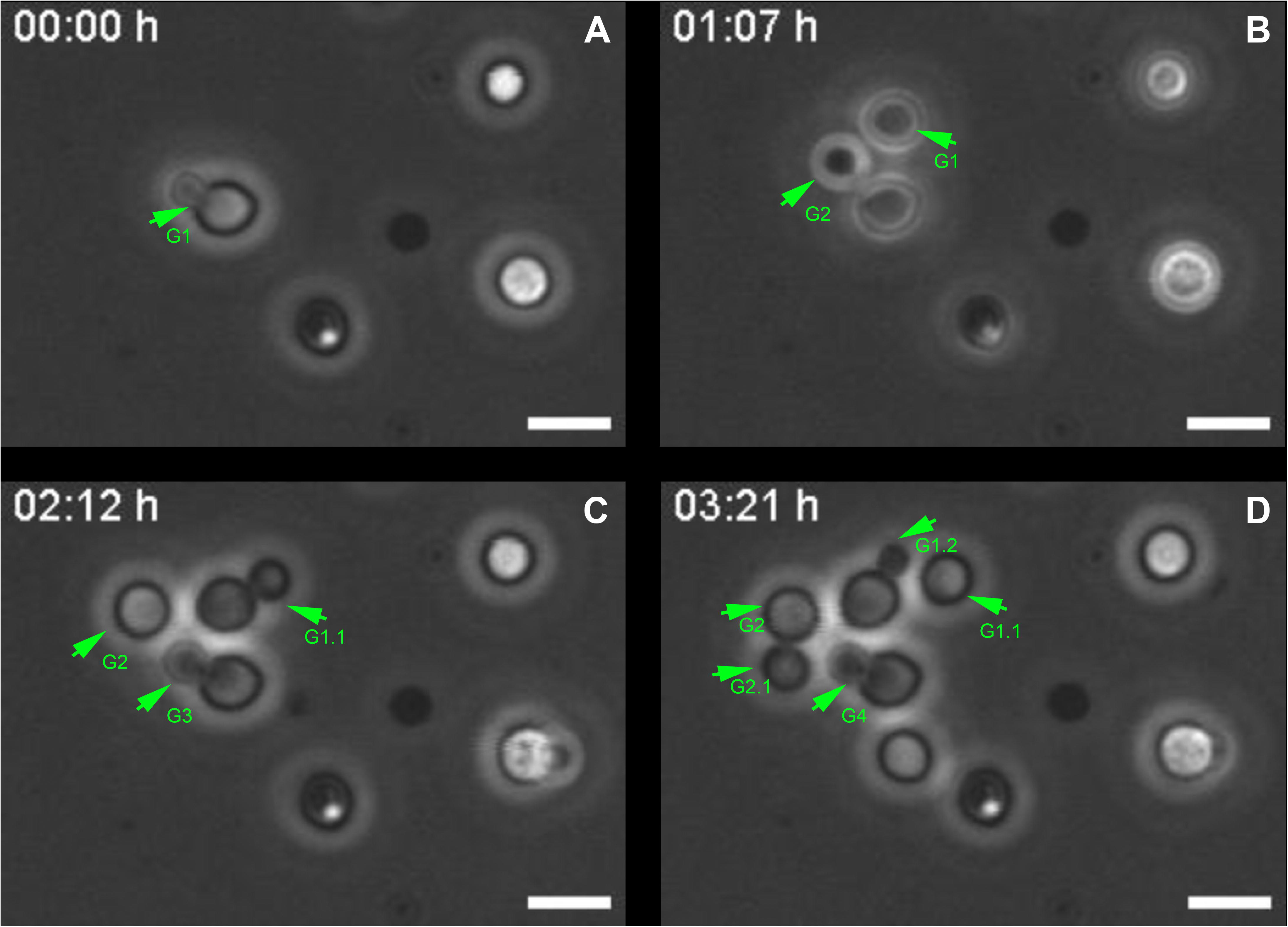
Selected images from a video microscopy movie of *C. neoformans*, grown in Sabouraud medium, showing budding evolution. A-D) Snapshots taken at 0, 1h07min, 02h12min and 03h21min, respectively, showing how the budding events evolve with time. Green arrowheads indicate yeasts during the division process that takes approximately (1.3 ± 0.3) h / cell. They also show that daughter cells always bud unipolarly and repeatedly from the same cell region. G1 indicates first generation cells, G1.1 daughter cell from the first generation, the other numbers follow a similar logic. Scale bar is 5 μm. (Supplementary video 1).

Although this observation is already a consensus in the *Cryptococcus* field (Zaragoza et al., 2006a, 2006b) it led us to hypothesize that the mother cell might develop a specialization at the cell wall, creating a region with certain characteristics that might facilitate budding.

### 3.2 *Cryptococcus neoformans* mother cell wall reorganizes prior to budding of daughter cells

In order to unravel details of these specialized regions (SRs), *Cryptococcus neoformans* yeast cell cultures were stained with Uvitex2B to label the chitin polymers present in the cell wall and subsequently were observed in a conventional fluorescence microscope. Images showed that mother cells, before budding of daughter cells, formed regions of lower fluorescence intensities when compared to the rest of the cell perimeter (Figures 2A, 2B and 2C).

**Figure 2:**
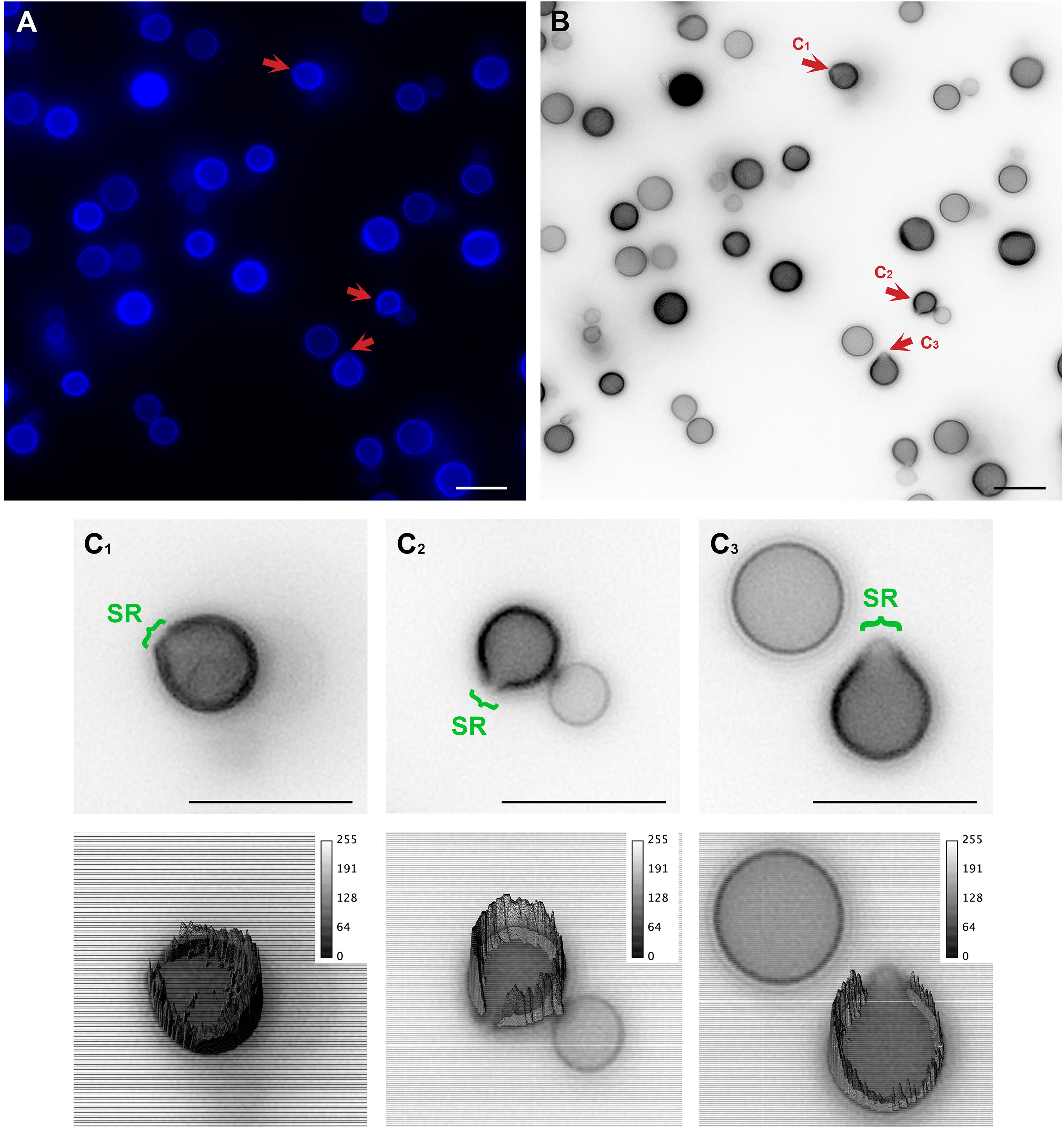
Conventional fluorescence microscopy images of *C. neoformans* stained with Uvitex2D depicting chitin polymers at the cell wall. **A)** Fixed and stained *C. neoformans* with chitin labeled in blue. Notice the discontinuity of the cell wall staining (red arrow). Those are the specialized regions (SR) in yeasts. **B)** Grey scale image of A. C1, C2 and C3 are cells that are under different budding stages (see red arrows). **C)** Zoom of C1, C2 and C3 (upper panel) with their respective 3D surface plots (lower panel). Surface plots indicate that the cell walls in SRs have a lower fluorescence intensity when compared to the entire cell wall rims. Scale bars are all 10 μm.

In an attempt to better clarify the organization of these SRs, we also imaged *C. neoformans* cells using SIM. Uvitex2B was used together with 18B7 + Alexa Fluor^®^ 594, to stain both the cell wall and the PS capsule respectively (Figure 3). The results confirmed the decrease in fluorescence intensities around the SR when compared to the entire cell perimeter (Figures 3A, 3B and 3C). Furthermore, although with few details, it was possible to identify that the cell walls seemed to peel off in order to form the SRs from which daughter cells bud. (Figure 3C and Supplementary Video 2).

**Figure 3:**
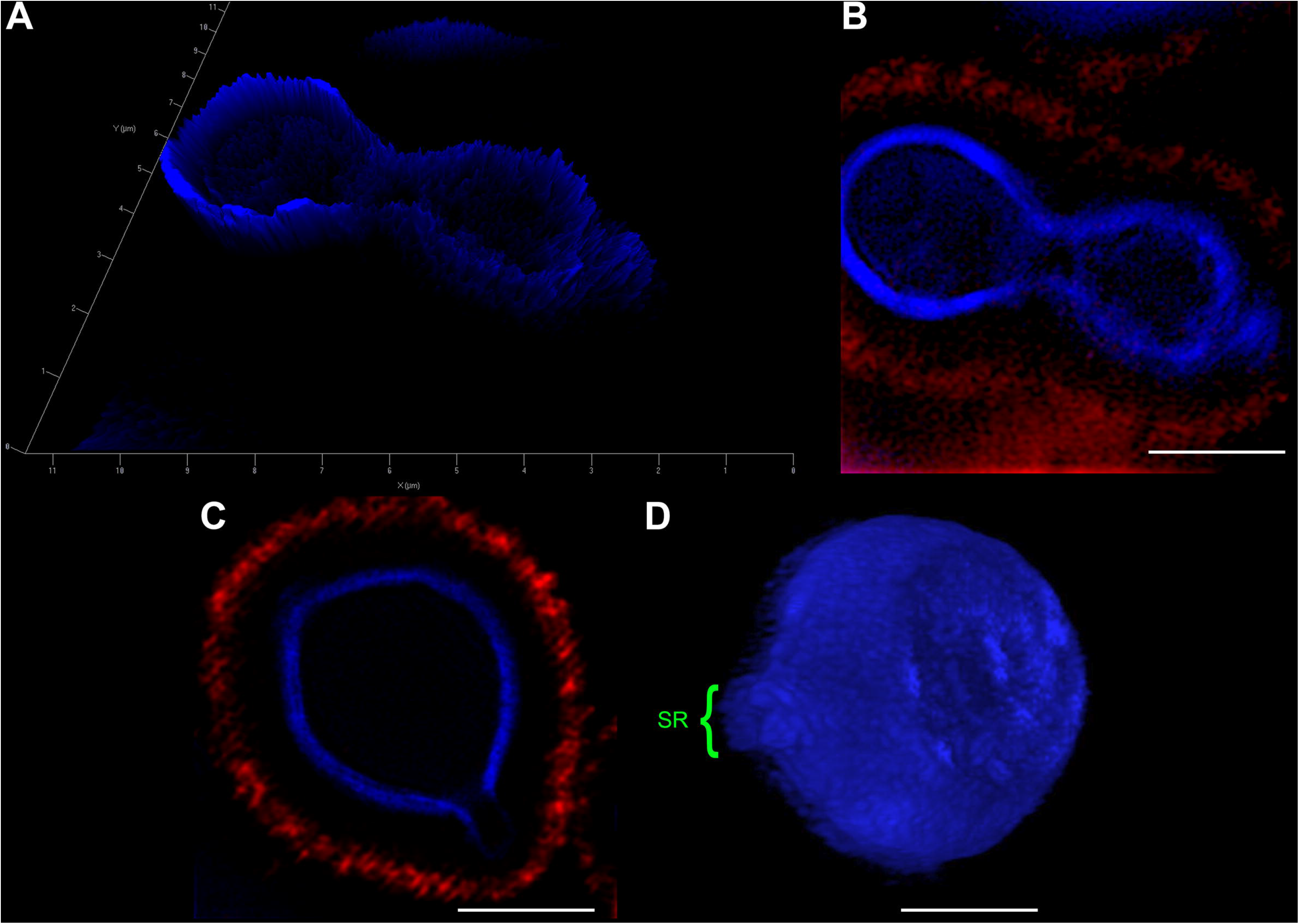
SIM images of *C. neoformans* confirm that both the cell wall and capsule deform prior to the budding event and generation of daughter cells. Yeast cells were fixed and stained for Uvitex2B (blue) and 18B7 antibody (red) so we could follow their walls and PS capsules, respectively, during budding and generation of daughter cells. **A)** 2.5D yeast profile, showing that the mother cell (left) has a higher cell wall fluorescence intensity when compared to the daughter cell (right). **B)** 2D representation of the same field as in A. **C)** Representative image of a yeast cell at the beginning of the budding process. SR can be visualized as a discontinuity of the cell wall. **D)** 3D profile of a representative yeast cell also in the beginning of the budding process showing the SR. Scale bars: A 1 μm. B, C and D 2 μm. (Supplementary video 2).

### 3.3 Ultrastructural details of *C. neoformans* specialized regions during budding

In order to better visualize the SRs with greater resolving power, we used transmission electron microscopy (TEM) and three-dimensional reconstruction by electron microscopy.

*Cryptococcus neoformans* yeast cell cultures were imaged using TEM. By using this technique, we were able to specifically follow the various steps of the budding process (Figures 4, 5 and 6). Our observations demonstrated that the process started with the shape change of one of the poles of the cell which deformed, lost sphericity and formed a more pointed region in the cell wall that, at some point, began to break into layers (Figures 4A and 4B). As the daughter cell started to appear, the formed layers became more evident with thick tips slightly separated from each other (Figures 4C and 4D). When the daughter cell is released, a new budding event is likely to arise in the same region as the first detached daughter cell turns the area more prone for subsequent occurrences (Figures 4E and 4F).

**Figure 4:**
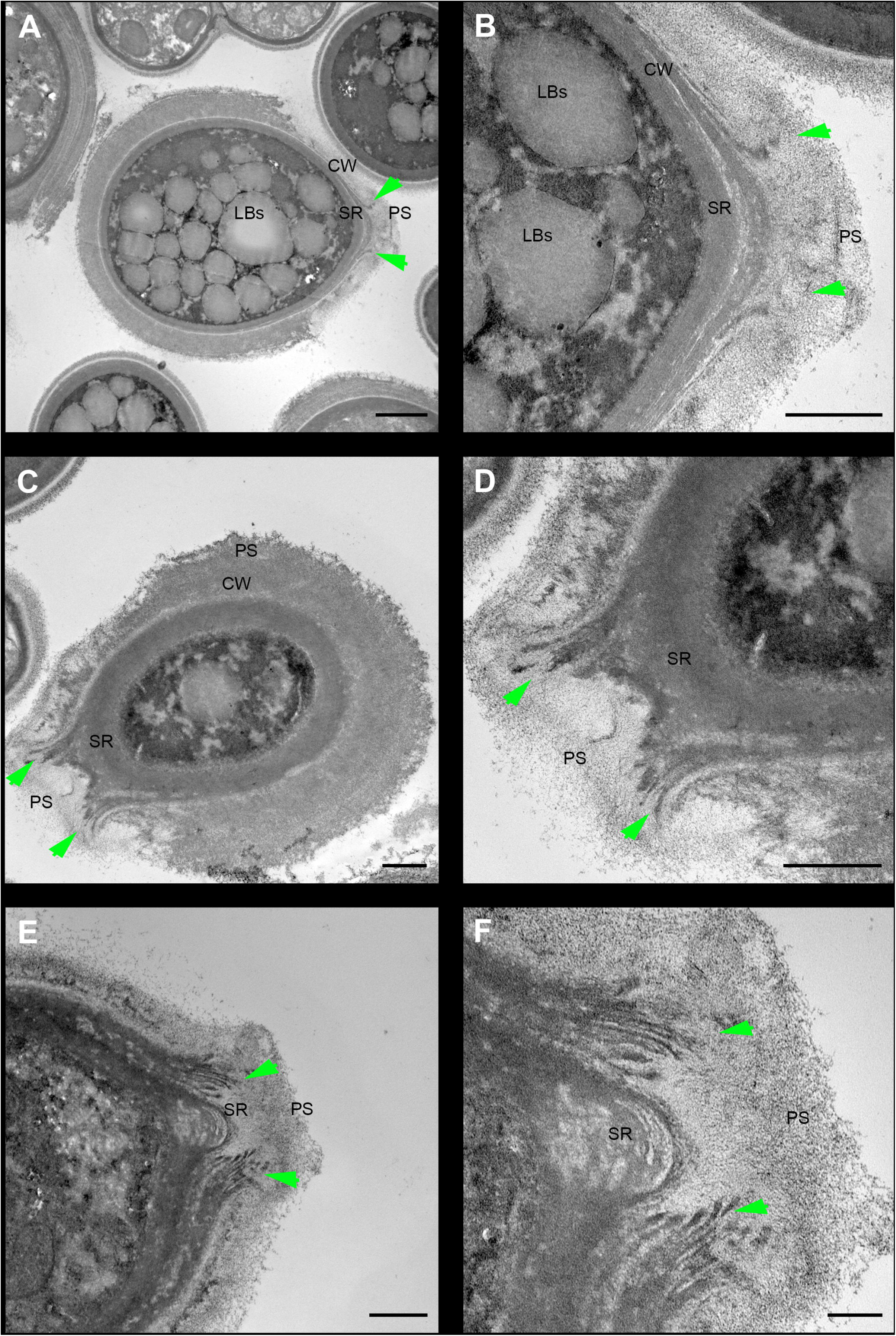
Transmission electron microscopy of *C. neoformans* during budding reveals that both the cell wall and PS layer reorganize prior to budding. The right column images (B, C, F) are zoomed images of the panels represented on the left. The green arrowheads indicate the cell wall delamination at different budding stages. **A - B)** Yeast in the beginning of the budding process presents a modest cell wall (CW) delamination in the specialized region (SR). **C - D)** Intermediate cell wall delamination stage and reduction of the PS fiber density at the SR when compared to the rest of the cell perimeter. **E - F)** Discontinuity of the cell wall showing an advanced delamination step in which the cell wall peeling is evident, resembling a “wafer cookie”, and the reduction of the density and complexity of the PS fibers around the SR is evident. **LBs:** Lipid Bodies; **CW:** Cell Wall; **PS:** Polysaccharide; **SR:** Specialized region. Scale bars: A, C and E 1 μm; B, D and F 500 nm.

Electron tomography and three-dimensional reconstruction were also performed in order to enable us to observe further details about the budding events (Figure 5 and Supplementary Video 3). Strikingly, with the tomography series, it became clearer that not only there is a separation of the mother cell wall into layers with thick tips (Figures 5A and 5B), as previously described (Figure 4), but also that the mother cell wall is thicker than the daughter cell wall (Figure 5D). Interestingly, daughter cells presented multilamellar membranous structures that seem to cover the continuous openings between daughter and mother cells, as a protective barrier (Figure 5D and 5E). It also became evident that the entire budding process induced capsule reorganization around the SRs (Figures 4 and 5). The most striking changes observed in the capsule morphology during budding were the reduction in PS density around SRs (Figure 4) and the formation of a protective PS barrier surrounding the SRs of both mother and daughter cells (Figures 5C, 5D and 4E). Three-dimensional reconstruction clearly demonstrated all of these characteristics (Figures 5F and 5G).

**Figure 5:**
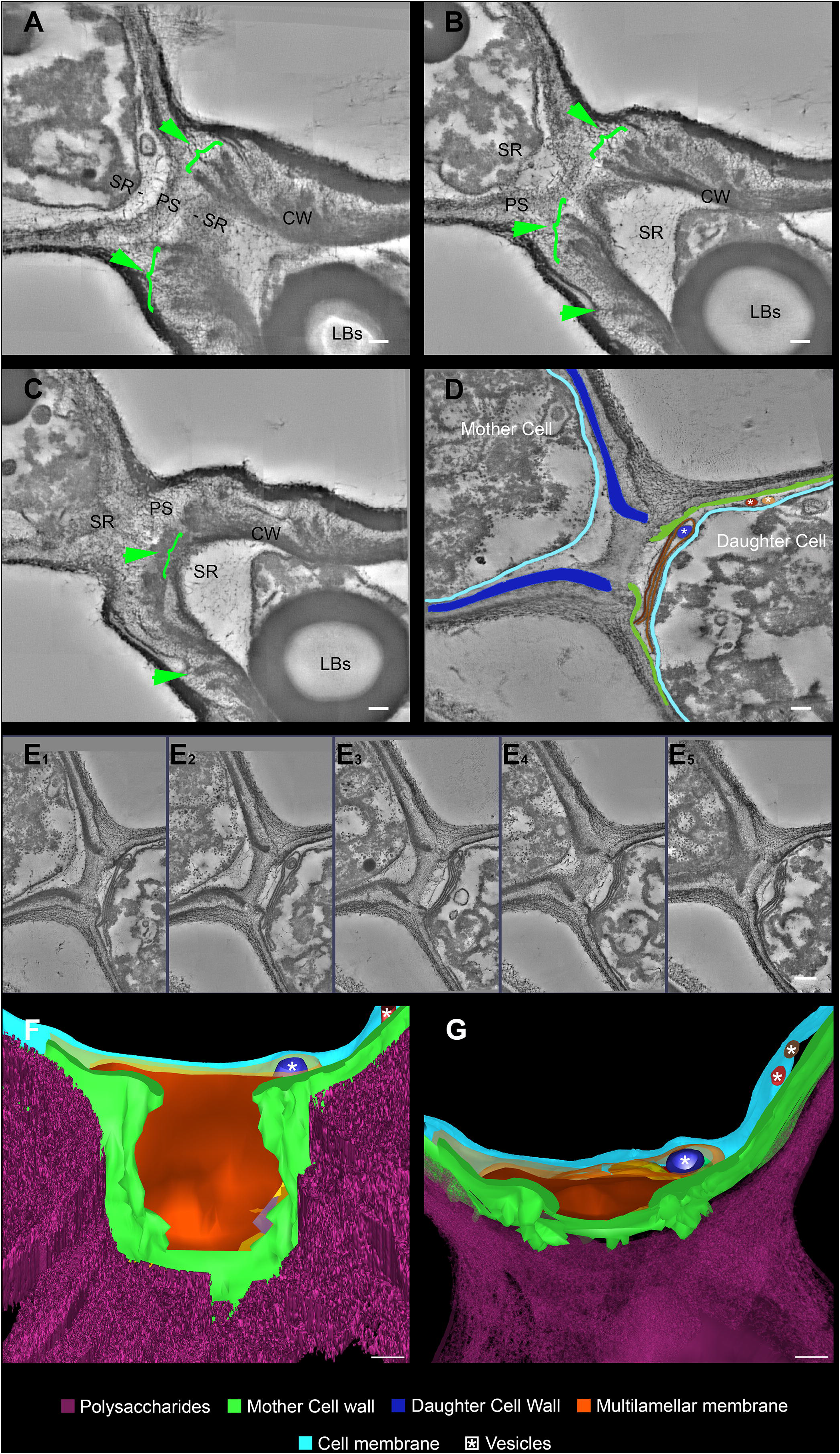
Serial electron tomography and three-dimensional reconstruction of *C. neoformans* show ultrastructural details of the budding process. **A – C)** Different z-planes of a serial tomogram of a budding event. The green arrowheads point to the cell wall delamination along different image planes. **D)** Virtual plane of the tomogram where the 3D model was superimposed to highlight the structures that participate and are remodeled in the SR. The upper left cell is the mother cell while the bottom right one is the daughter cell. **E_1-5_)** Set of sequential slices from the tomogram evidencing the modifications in cell wall cohesion along different angles. Note the presence of a multilamellar membrane and vesicles in synergism with the PSs surrounding the SRs, which probably acts as a shield. **F and G)** Different views of the three-dimensional model from the daughter cell perspective. In **F** we see a front view of the daughter cell wall discontinuity at the region of budding. This region is sealed by several membrane profiles as show in E. In G, a top view of the model where membrane profiles can be seen in between the CW and the cell membrane. Several vesicles (asterisk) could be seen inside the membrane profiles (dark blue) and also between CW and cell membrane (red and brown). **LBs:** Lipid Bodies; **CW:** Cell Wall; **PS:** Polysaccharide; **SR:** Specialized region. Scale bars are all 100 nm. (Supplementary video 3).

Thus, cell wall structural reorganization and capsule remodeling are evident processes that occur in both mother and daughter cells during budding.

### 3.4 Capsule polysaccharide fibers around specialized regions of daughter cells are oriented towards the budding events

In order to clarify the details of capsule remodeling and also to better understand the PS reduced density around the SRs, *C. neoformans* were processed and visualized by conventional and high-resolution scanning electron microscopes (Figure 6).

**Figure 6:**
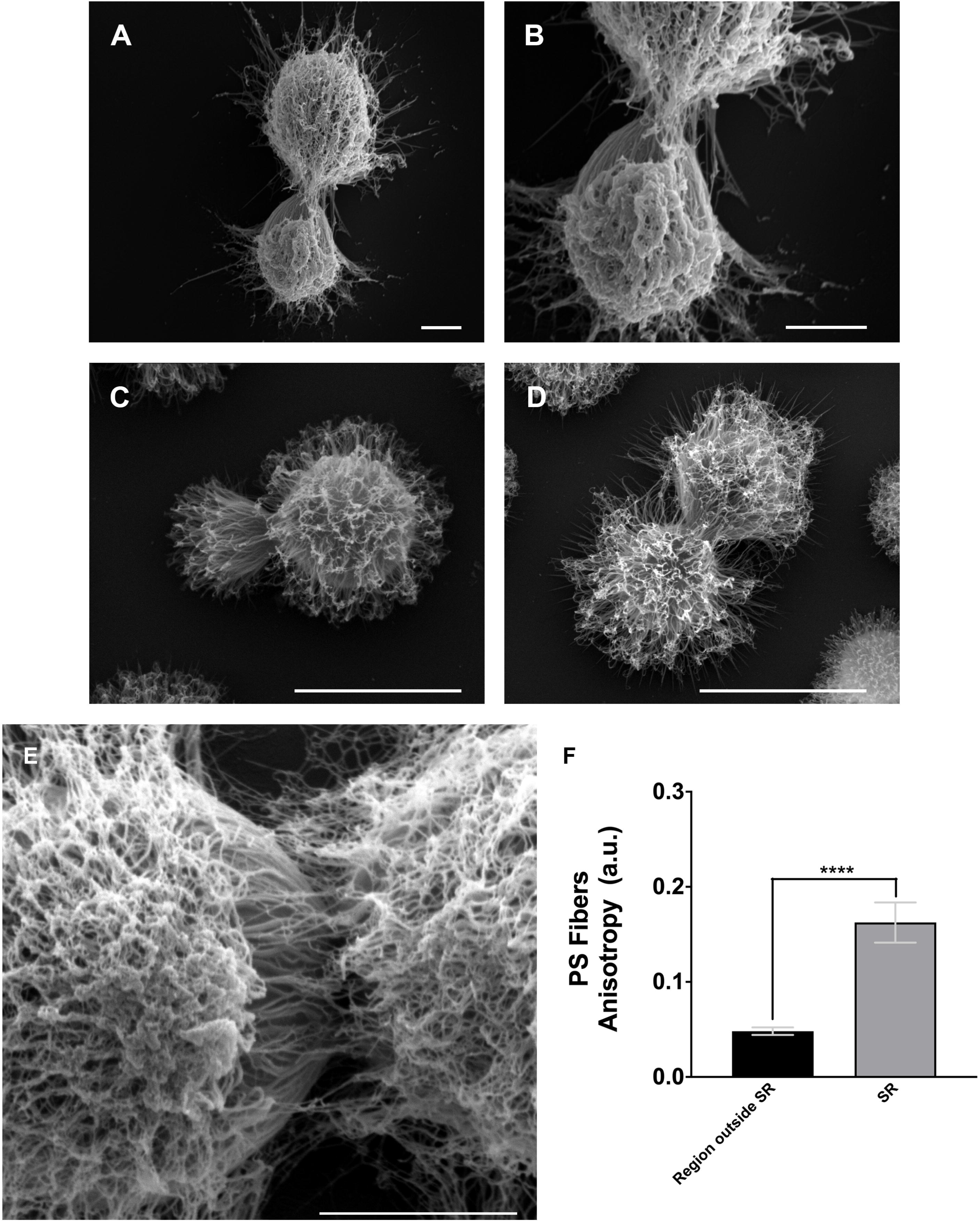
Scanning electron microscopy images of *C. neoformans* showing the alignment of the PS fibers toward the budding region. **A - E)** Different examples showing changes in the spatial and conformational orientation of the PS capsule in the SR. Scale bars: A, B and E 2 μm; C and D 5μm. **F)** Plot of the mean anisotropy values of PS fibers around SRs (grey) and outside SRs (black). At least 25 different measurements were performed for each experimental situation. Standard error was used as error bars. **** means *p* < 0.0001 in Student’s *t*-test statistics.

As previously described (Figures 4 and 5), the images presented in Figure 6 confirmed not only the remodeling but also the reduction in PS density around the SRs. Moreover, the PS fibers from daughter cells have a distinct spatial organization surrounding these SRs, with fibers aligned in the direction of budding (Figures 6A – 6E). In order to better quantify these visual observations, we performed a quantitative analysis using the FibrilTool plug-in (Boudaoud et al., 2014). The results (Figure 6F) showed that, overall, the PS fibers around SRs present a higher anisotropy when compared to fibers outside the SRs. This special arrangement around SRs may be related to the mechanical force that occurs during budding.

## 4. Discussion

*Cryptococcus neoformans* present themselves as haploid and spherical yeasts, surrounded by a PS capsule, unique feature among eukaryotes (D. C. McFadden et al., 2006). PS capsule in *C. neoformans* is its main virulence factor (Doering, 2000; Rodrigues et al., 2009; Zaragoza, 2019; Zaragoza et al., 2009), and it plays several roles, such as protection against dehydration and phagocytosis by natural predators in the environment. In addition, in the context of human infection, the PS capsule provides protection against phagocytosis, inhibition of leukocyte migration, depletion of the complement system and inhibition of antibody production (Dong et al., 1995; Feldmesser et al., 2001; Kozel et al., 1977; Macher et al., 1978; Murphy and Cozad, 1972; Retini et al., 1998; Zaragoza et al., 2009, 2008). Thus, the PS capsule becomes a primary component of interaction between the fungi and their host cells, using its immunomodulatory dynamics and its antiphagocytic properties to hinder the host’s immune response, therefore becoming one of the main targets for therapeutic strategies (Casadevall and Pirofski, 2007; Larsen et al., 2005). In addition, and just underneath the PS capsule, there is a rigid structure called the cell wall. It is considered to be the primary determinant of *C. neoformans* resistance to stress and environmental aggressions. The cell wall architecture, thoroughly described in the introduction of the present study, consists mainly of a polymer network, including chitin, glucans, manans and galactomanans as its major components (Roncero, 2002; Ruiz-Herrera et al., 2002).

Polarized cell growth (PCG) and directional cell division (DCD) are fundamental and essential processes for the development of eukaryotes. PCG involves asymmetric growth of a cell region to form specific cell structures or shapes. The resulting specialized structures are critical for the function of several cell types and can help mediate various cell interactions during development. Some examples are the absorption of nutrients by epithelial cell microvilli (Mooseker, 1985) and the interaction between T and B cells (Kupfer et al., 1986; Madden and Snyder, 1998). Likewise, PCG in fungi occurs by inserting new material into the plasma membrane via the secretory pathways together with concomitant cell wall remodeling. It can be either triggered by internal factors, for example progression of the cell cycle, or by external factors, such as changes in the environment or nutritional status (Bassilana et al., 2020). Although filamentous fungi and yeasts show obvious differences in their growth modes, they share three basic properties that allow for PCG and the formation of a diverse variety of cellular forms: **(I)** symmetry breaking, in which an initially isotropic cell generates a polarized growth axis, **(II)** maintenance of polarity, which refers to the stabilization of the polarity axis, so that polar growth is maintained, and finally **(III)** depolarization, in which polarity is lost in a controlled manner. The balance between polarity maintenance and depolarization generates the diversity in fungal cell forms (Lin et al., 2014). Fungal cells are not always polarized during the early stages of development. They usually undergo an initial period of non-polar isotropic expansion (for example, spores from yeast stem cells). Ultimately, however, cell symmetry must be broken, and a polarity axis generated, both for the selection of the sprouting site or for the development of polar structures, such as hyphae.

Many of the studies that focused on cell division events for pathogenic yeasts used paradigms established in the ascomycete *Saccharomyces cerevisiae*. However, basidiomycete yeasts, such as *C. neoformans*, show conserved and distinct features in their morphogenesis. It only produces hyphae during sexual differentiation (Lin et al., 2014) and their produced spores are quite infectious and can be the primary particle inhaled during a natural infection (Giles et al., 2009; Velagapudi et al., 2009). Once inhaled, upon reaching the lungs, the Cryptococci spores germinate to produce yeast cells. In the context of infection, this fungus grows within the human host almost exclusively in the form of yeasts. Histopathological studies have shown that hyphal forms are rarely found during human infections by *C. neoformans*. and elongated fungal morphologies are only observed as rare variants among clinical strains (Baker and Haugen, 1955; Fu et al., 2019; Shadomy and Utz, 1966).

Cellular events during *C. neoformans* yeast cell morphogenesis are well described, but still lack information regarding ultrastructural changes. In general, cells undergo a cell division cycle with asexual and repeated clonal budding of a haploid yeast cell; however, unlike *S. cerevisiae*, in which subsequent sprouting events occur adjacent to previous scars, *C. neoformans* preferentially and repeatedly generates their daughter cells from the same location, as shown in the present study. For this reason, scar count, as a result of budding, is unlikely to provide an accurate age measurement of cryptococcal cells (Adams, 2004; Zhao et al., 2019).

The fungal cell wall is a protective barrier that resists environmental and osmotic stresses, maintaining cell morphology and regulating membrane permeability, as well as a wide range of essential roles during the fungus’s interaction with its environment (Free, 2013; Gooday, 1995). However, in spite of serving as a protective structure, the cell wall also needs to be remodeled and even partially ruptured for the subsequent budding events that will allow the perpetuation of the fungus in its chosen infection site. Our results not only corroborate this hypothesis but also describe for the first time, to the best of our knowledge, the morphological features of how this remodeling occurs. We showed that the cell wall starts to form a fissure-like structure that culminates with the opening of a specialized region where *C. neoformans* preferentially and repeatedly generates its daughter cells. The cell wall is then reorganized into layers similar to “wafer cookies” around the specialized region. We conjecture that the formation of this region is related to a protruding force coming from inside the cell.

However, apart from the mechanical aspect, we cannot completely rule out the possibility of a change in biochemical composition to facilitate this process. It is known that the cell wall is divided into two layers with specific components. The inner layer is mainly composed of β-glucan and chitin arranged as fibers parallel to the plasma membrane and the outer layer contains α-glucan and β-glucan (O’Meara and Alspaugh, 2012; Sakaguchi et al., 1993). Moreover, several cell wall proteins have been described to have key roles in the capsule architecture. Some of these proteins, such as the GPI-linked β-glucanase Gas1, have been implicated in remodeling of the cell wall as it directly acts on β-1,3-glucans (Eigenheer et al., 2007; Levitz and Specht, 2006). Therefore, preceding cell budding, perhaps still during the PCG stage, the structural components of the cell wall may be reorganized in order to form a region where budding is facilitated. In the present study, we have described the morphological aspects of the cell wall remodeling. The molecular and mechanical details constitute lines of investigation for future studies.

Beyond the cell wall and attached to its surface lies the PS capsule. Despite its homogeneous appearance, when viewed through light microscopy, several lines of evidence show that the capsule is a highly heterogeneous structure with a complex and dynamic spatial organization. It is known, for example, that the capsule matrix exhibits clear vertical stratification, with distinct density regions, with its inner part having a higher fiber density than its outer region (Araújo et al., 2016, 2017; Bryan et al., 2005; S. Frases et al., 2009; Gates et al., 2004). Although softer than the cell wall and presenting viscoelastic behavior (Araújo et al., 2019; Susana Frases et al., 2009), the high PS density of the inner region prevents the penetration of larger macromolecules, including antibodies and proteins of the complement system, restricting the access of these molecules to the cell wall (Gates et al., 2004; Gates and Kozel, 2006). Therefore, one could speculate that the capsule might also undergo remodeling during budding. Indeed, our results show that the PSs constituting the capsule undergo shape changes around the specialized budding region. The change in shape seems to be correlated with the conjectured protruding force, as the capsule fibers tend to orient towards the budding event. Finally, our results also demonstrate that the PS capsule works as a protective shield around the specialized region during the budding event.

In conclusion, we have combined the state-of-the-art in light and electron microscopy techniques to describe the structural changes that synergistically occur between the capsule, cell wall and plasma membrane during the budding phenomenon in *C. neoformans*. We have presented evidences supporting a possible remodeling through mechanical protruding forces originated inside the yeast cell, although the magnitude of such force and the mechanism behind its generation have yet to be elucidated. All morphological changes observed, particularly those from the cell wall, act together and create a specialized region with characteristics that seem to favor budding events, which is able to partially explain why budding in *C. neoformans* always occur in the same region. However, the complete mechanism may also involve controlled rearrangement of the molecules that constitute both the cell wall and the PS capsule and will be explored in future studies. We also aim to explore and characterize further these specialized regions such that they could be used as potential drug-targets against cryptococcosis.

## 5. Author Contributions

**Glauber R. de S. Araújo:** Conceptualization, Methodology, Investigation, Visualization, Data curation, Formal analysis, Writing - original draft, Writing - review & editing.

**Carolina de L. Alcantara:** Methodology, Writing - review & editing

**Noêmia Rodrigues:** Methodology, Writing - review & editing

**Wanderley de Souza:** Supervision, Funding acquisition, Formal analysis, Writing - review & editing

**Bruno Pontes:** Conceptualization, Methodology, Investigation, Visualization, Data curation, Resources, Supervision, Funding acquisition, Formal analysis, Writing - original draft, Writing - review & editing.

**Susana Frases:** Conceptualization, Methodology, Investigation, Visualization, Data curation, Resources, Supervision, Funding acquisition, Formal analysis, Writing - original draft, Writing - review & editing.

## 7. Acknowledgments

We acknowledge Dr. Barbara Hissa for critical reading and scientific editing of the manuscript. We also thank the technicians from Centro Nacional de Biologia Estrutural e Bioimagem (CENABIO/UFRJ) for all-important help. This work was supported by the Brazilian agencies Conselho Nacional de Desenvolvimento Científico e Tecnológico (CNPq), Coordenação de Aperfeiçoamento de Pessoal de Nível Superior (CAPES) - Finance Code 001 and Fundação Carlos Chagas Filho de Amparo à Pesquisa do Estado do Rio de Janeiro (FAPERJ).

## 8. Appendix A. Supplementary data

Supplementary video 1, Supplementary video 2 and Supplementary video 3.

## 10. Supplementary Material Legends

**Supplementary video 1:** Budding phenomena observed in a representative *C. neoformans* cell culture (related to Figure 1).

**Supplementary video 2:** 3D-SIM reconstruction of a representative *C. neoformans* cell in the beginning of the budding process showing the SR (related to Figure 3D).

**Supplementary video 3:** Set of electron tomography planes showing *C. neoformans* mother and daughter cells during a budding process (related to Figure 5E).

